# Transcriptomic analysis of loss of Gli1 in neural stem cells responding to demyelination in the mouse brain

**DOI:** 10.1101/2021.02.28.433246

**Authors:** Jayshree Samanta, Hernandez Moura Silva, Juan J. Lafaille, James L. Salzer

## Abstract

In the adult mammalian brain, Gli1 expressing neural stem cells reside in the subventricular zone and their progeny are recruited to sites of demyelination in the white matter where they generate new oligodendrocytes, the myelin forming cells. Remarkably, genetic loss or pharmacologic inhibition of Gli1 enhances the efficacy of remyelination by these neural stem cells. To understand the molecular mechanisms involved, we performed a transcriptomic analysis of this Gli1-pool of neural stem cells. We compared murine NSCs with either intact or deficient Gli1 expression from adult mice on a control diet or on a cuprizone diet which induces widespread demyelination. These data will be a valuable resource for identifying therapeutic targets for enhancing remyelination in demyelinating diseases like multiple sclerosis.

## BACKGROUND & SUMMARY

Multiple Sclerosis (MS) is the most common cause of neurological disability in young adults^1^ and is characterized by inflammatory demyelination leading to axonal injury. In addition to preventing immune-mediated demyelination, a major therapeutic goal in MS is to enhance new myelin sheath formation by remyelination which prevents the loss of axons and restores function^2–4^. The nervous system is capable of remyelination as observed in autopsy specimens from MS patients characterized by short, thin myelin sheaths^5–8^. However, remyelination ultimately fails, particularly in MS patients with progressive disease^8,9^ and the factors that limit remyelination remain poorly understood^9,10^.

There are three sources of remyelinating cells in the adult brain - neural stem cells (NSCs)^11–13^, oligodendrocyte progenitor cells (OPCs)^14–16^ and mature oligodendrocytes^17,18^. NSCs reside in the subventricular zone (SVZ) while the OPCs and oligodendrocytes are present throughout the parenchyma in the adult brain and are recruited locally to regenerate myelin in demyelinated lesions^15,19^. Furthermore, NSCs can promote remyelination either by replenishing the adult OPCs recruited into lesions, or as a direct source of oligodendrocytes themselves. Consistently, prior studies have shown that NSCs in the SVZ are activated by local demyelination, migrate out and differentiate into OPCs that generate remyelinating oligodendrocytes^11,13,14,20^. Indirect marker studies of postmortem brains further suggest that NSCs proliferate during acute attacks of MS and are a source of remyelinating cells in the human brain^12,21,22^. More importantly, ablating NSCs results in axonal loss highlighting their significant functional contribution to remyelination^23^. Taken together, these studies suggest remyelination by NSCs protects axons. Thus, enhancing remyelination by NSCs is expected to provide a neuroprotective strategy in MS.

The adult SVZ is comprised of a functional, quiescent neural stem cell population, which are multipotent, divide slowly, self-renew, and give rise to proliferating, transit-amplifying cells^24^. There is considerable heterogeneity among these NSCs, indicated by their expression of specific transcription factors and generation of distinct cells during homeostasis and in response to injury^25^. One NSC subset, which is enriched in the ventral SVZ, expresses the transcription factor Gli1 and generates interneurons in the olfactory bulb and astrocytes in the healthy mouse brain^26,27^. This population of NSCs is also present in the human SVZ^27^ as well as in NSCs derived from human induced pluripotent stem cells and human embryonic stem cells^28^. Remarkably, and in contrast to its fate in the healthy adult brain, this Gli1 pool of NSCs is specifically recruited to the white matter in response to demyelination and goes on to differentiate into remyelinating oligodendrocytes^20,27,29^. Inhibition of Gli1 in these cells further enhances their remyelination potential and is neuroprotective resulting in functional improvement in mouse models of MS^27,28^. Importantly, Gli1 expressing stem-like cells respond to injury in other organs like the liver, lung, bone marrow, kidney and heart with loss of Gli1 resulting in attenuated injury in these organs^30–37^. Thus, inhibition of Gli1 is potentially an important therapeutic strategy for regenerative medicine and the molecular mechanisms by which Gli1 affects stem cell repair may be shared in many different types of tissues.

We performed a transcriptomic analysis to identify Gli1-regulated genes that affect NSC recruitment and generation of remyelinating oligodendrocytes by comparing the mRNA expression between NSCs with genetic loss of Gli1 and those with intact Gli1 expression from both healthy and demyelinated adult mouse brains (Fig. 1). These data will be valuable not only for future remyelination studies but may also provide mechanistic insights into the shared principles of regeneration across organs.

**Figure 1:**
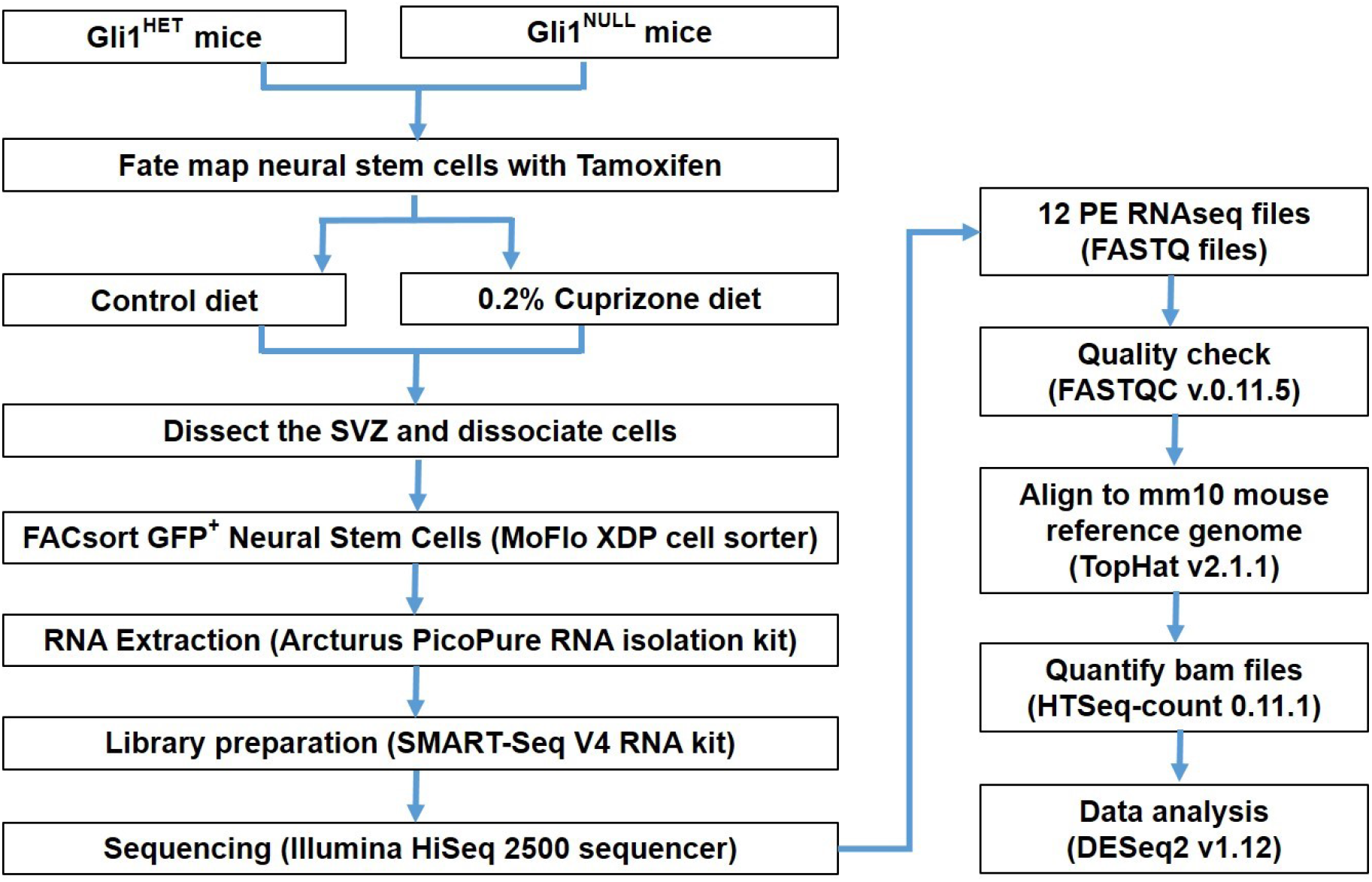
Schematic overview of the study design. A flowchart demonstrates the experimental design and data analysis.

## METHODS

### Fate-mapping of neural stem cells and demyelination

All the mice had access to food and water *ad libitum* and were housed in a room with 12 hour dark/light cycle. The genetically modified mice belonged to C57bl/6 strain and were maintained according to protocols approved by New York University Medical Center’s IACUC. The *Gli1^CreERT2/+^* mice (Jax # 007913) which have CreERT2 knocked into the Gli1 locus were bred with the reporter Rosa-CAG-EGFP-LoxP (RCE) (Jax# 032037) mice to generate *Gli1^HET^* (*Gli1^CreERT2/+^;RCE*) with intact *Gli1* expression and *Gli1^NULL^* (*Gli1^CreERT2/CreERT2^;RCE*) mice with global knockout of *Gli1*. To permanently label the Gli1 pool of NSCs with GFP, we administered 5 mg tamoxifen (Sigma) in corn oil on alternate days for a total of four intraperitoneal injections to 10 week old mice^27,38^. We did not observe any labeling in the absence of tamoxifen administration. Mice used for GFP negative controls received intraperitoneal injections of corn oil without Tamoxifen. A week after the last dose of tamoxifen/corn oil, the demyelinated group was fed 0.2%cuprizone in the chow for 3 weeks while the healthy control group remained on regular chow. Cuprizone diet induces apoptosis of mature oligodendrocytes leading to maximum demyelination of the white matter corpus callosum after 5 weeks of continuous feeding^39^. Here, we used 3 weeks of cuprizone diet for our analysis since we previously found that *Gli1^NULL^* NSCs in the SVZ have significantly higher proliferation at this timepoint compared to *Gli1^HET^* SVZ, suggesting that maximal activation of NSCs coincides with this early stage of demyelination^27,40^.

### Dissection of SVZ

Brain tissues were harvested from 6 mice (3 Males and 3 Females) per genotype (*Gli1^HET^* and *Gli1^NULL^*) per diet (cuprizone and control) after euthanizing them with CO_2_. The brain was removed gently out of the skull and immediately transferred to a Petri dish chilled on ice and placed with the dorsal surface facing up on a chilled acrylic mouse brain matrix. The brains of mice were dissected according to previously published protocols^38^. Coronal sections of 1 mm thickness were made from the rostral end of the brain up to the dorsal hippocampus posteriorly, using a sharp and chilled razor blade. The SVZ was identified as a thin layer of tissue lining the medial and lateral walls of the lateral ventricles and carefully dissected under a microscope from each coronal brain section.

### Dissociation and FACsorting of neural stem cells

For each RNA sample, the SVZ from one female and one male mice was pooled together. Overall, we had 3 RNA samples/genotype from control diet group and 3 RNA samples/genotype from cuprizone diet group in this study. The SVZ was then dissociated into single cell suspension following the manufacturer’s protocol in the papain dissociation kit (Worthington Biochemicals # LK003150)^41^. Briefly, the dissected SVZ tissue was minced using iris scissors and digested by incubating in a solution containing 20 Units/ml papain and 0.005% DNase for 30minutes at 37°C. Subsequently, the papain was inactivated with albumin-ovomucoid inhibitor and the digested tissue was needle triturated to obtain a cell suspension^41^.

The cell suspension was passed through a 50 μm celltrix filter (Sysmex-Partec # 04-0042-2317) immediately before FACsorting the GFP+ NSCs using a MoFlo XDP cell sorter (Beckman Coulter) in the NYU Cytometry core facility. NSCs harvested from *Gli1^NULL^* mice treated with corn oil served as negative controls for FACsorting. A forward scatter vs. side scatter dot plot was first used to gate the primary cell population and eliminate dead cells and debris. The gating was further refined to exclude doublets to finally select GFP+ cells (Fig. 2. Gate strategy exemplified by magenta arrows). In order to minimize any perturbation of gene expression, we performed cell sorting based on cell size and GFP expression immediately after obtaining a single cell suspension without manipulating the cells. In order to prevent the perturbation of gene expression, we did not add a viability dye to the cells during the FACsorting. However, we used 10 μl of the sorted GFP+ cells in phosphate buffered saline (PBS) for trypan blue cell viability assay and quantified the clear viable cells along with the blue stained dead cells using a hemocytometer^42^. All our FACsorted samples consisted of more than 95% viable cells. On average, the cell sorting yielded 17,666.7 ± 4163.3 GFP+ cells in the *Gli1^HET^* healthy group on control diet, 7,833.3 ± 2,112.7 GFP+ cells in the *Gli1^HET^* demyelination group on cuprizone diet, 41,333.3 ± 5033.2 GFP+ cells in the *Gli1^NULL^* healthy group on control diet and 34,000 ± 10,816.7 GFP+ cells in the *Gli1^NULL^* demyelination group on cuprizone diet.

**Figure 2:**
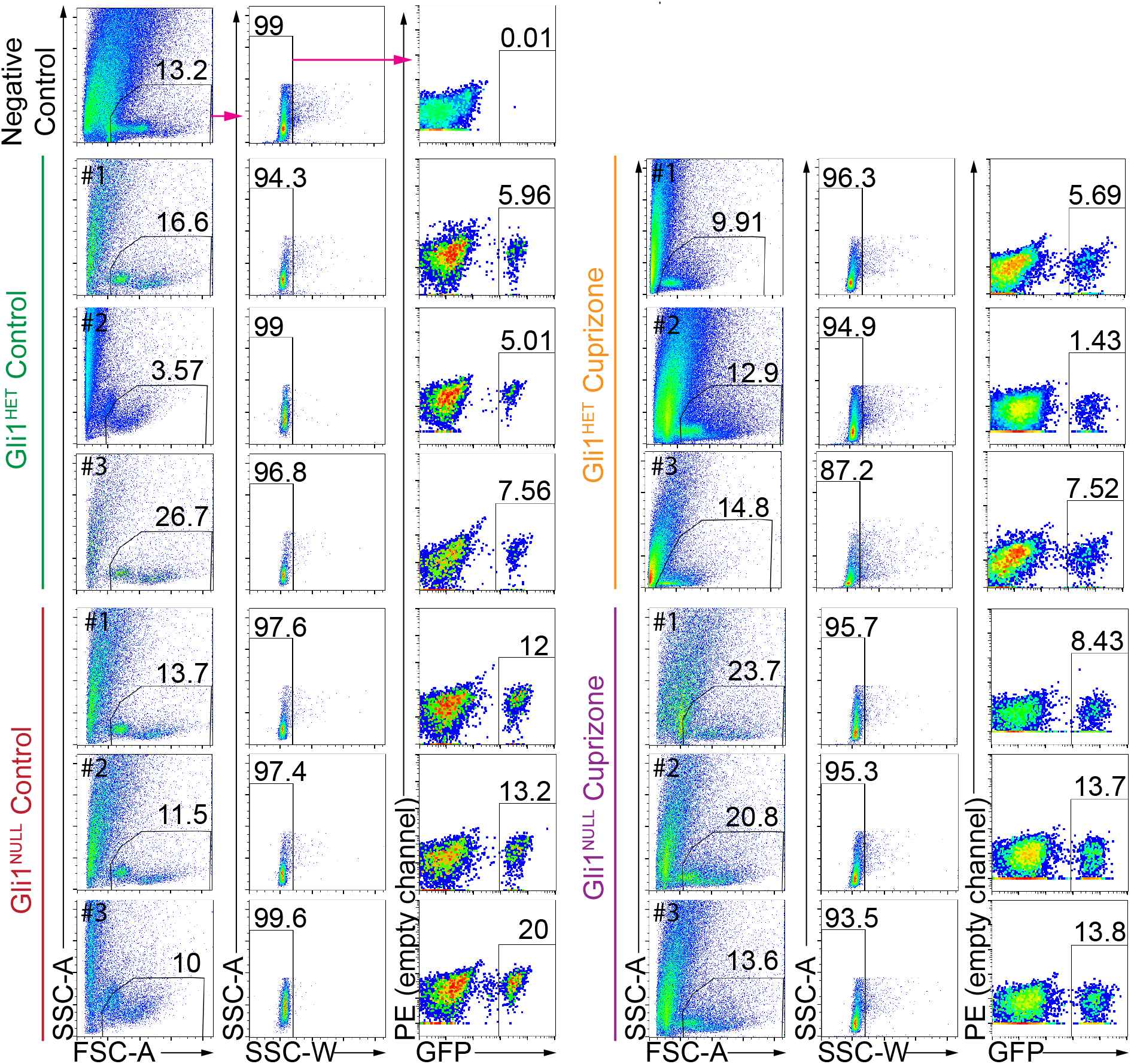
Flow cytometry analysis displaying the gating strategy used to sort fate-mapped neural stem cells. The negative control (top row) displays the gating strategy highlighted by the magenta arrows. The same strategy was applied for all samples in the figure. Fate-mapped, (i.e. GFP+) neural stem cells were purified from each of the samples depicted in the figure and processed for RNAseq. Percentage of gated cells is displayed above the selected area and the sample number is indicated in the top left corner of each dot plot. SSC-Side scatter, FSC-Forward scatter.

### Total RNA isolation

The GFP positive neural stem cells were collected in RNA extraction buffer (Arcturus PicoPure RNA isolation kit, Applied Biosystems # 12204-01) after FACsorting and total RNA was isolated following the manufacturer’s instructions. The isolated RNA was eluted in 15 μl of elution buffer following on column DNAse1 treatment using RNase-free DNase set (Qiagen # 79254). The total RNA was assessed for RNA integrity with a bioanalyzer (Agilent) in the NYU Genome Technology Center. RNA samples with a RIN of more than 8 were submitted to NYU Genome Technology Center for Next Generation Sequencing (NGS).

### Library preparation and RNA sequencing

mRNA was purified and the cDNA was synthesized with the mRNA fragments as templates using SMART-Seq V4 RNA kit (Clontech # 634889). The sequencing libraries were generated using Nextera XT DNA Library Preparation Kit (Illumina #FC-131-1024), which uses an engineered transposome to simultaneously fragment and barcode the cDNA, adding unique short adapter sequences. The barcoded cDNAs were then amplified by 6 cycles of PCR and their quality was checked for size, quantity and integrity by TapeStation system (Agilent) as well as qPCR for precise quantification. The generated libraries were then mixed for multiplexing on 3 lanes and sequenced on Illumina HiSeq 2500 platform with 150bp paired end modules, which yielded mapped reads in the range of 16 - 39 million per sample.

### Data transformation and downstream analysis

The data were transformed and analyzed by a bioinformatician in the NYU Genome Technology Center. Briefly, the raw sequencing data files were initially processed using FASTQC v.0.11.5 in order to perform quality control checks. Subsequently the Fastq files were aligned to the mm10 mouse reference genome utilizing TopHat v2.1.1. About 93% to 96% of the RNAseq reads, including R1 reads in the 5’ to 3’ forward direction and R2 reads in the reverse direction, mapped to the mouse genome using the Ensembl gene annotation system^43^. The aligned bam files were then quantified for gene expression values utilizing HTSeq-count 0.11.1. Finally DESeq2 v1.12 was used for analysis of the quantified count matrix^44^. Genes that had an adjusted p-value (False Discovery Rate) of less than 0.05 were considered significantly different for respective pairwise comparisons.

## DATA RECORDS

The sequencing data have been deposited in NCBI’s Gene Expression Omnibus and are accessible through GEO Series accession number GSE162683^45^. Table 2 lists the accession number for each sample. The data related to RNA quality, barcode lane statistics, RNA read counts, alignment statistics, DEseq2 expression counts and the data comparison between the groups have been deposited in Figshare^46^.

## TECHNICAL VALIDATION

### FACsorting

Since *Gli1^NULL^* mice carry two copies of the *CreERT2* allele, they would be expected to have a higher probability of tamoxifen independent leaky GFP expression. Hence, we used cells isolated from the brains of *Gli1^NULL^* mice but injected with corn oil instead of tamoxifen such that GFP is not expressed, as our negative control for FACsorting. These GFP negative cells were used to set up the gating parameters before GFP positive cells were sorted which ensured that only GFP positive neural stem cells were collected after sorting (Fig. 2).

### RNA quality

The quality of total RNA was assessed by RNA Integrity Number (RIN) using the Agilent Bioanalyzer. Following the standard practice in NGS, we used RNA with RIN>8 for our sequencing analysis^47^.

### Genotypic and phenotypic assessment

Before harvesting the neural stem cells, we collected the tail tissue from all the mice following euthanasia. These tissues were used for PCR analysis to reconfirm the genotypes of the mice used to isolate neural stem cells. We also obtained a higher number of GFP positive cells from *Gli1^Null^* mice compared to *Gli1^Het^* mice, consistent with higher proliferation rates of NSCs in the *Gli1^Null^* mice^27^, further validating their phenotype. An analysis of differential gene expression in healthy *Gli1^Het^* NSCs vs. healthy *Gli1^Null^* NSCs showed a significant difference in the expression on only one gene i.e. *Gli1*, which not only confirmed the genotype but also confirmed the well-established lack of phenotype in the healthy *Gli1^Null^* mice^48^ (Fig. 3 and Table 1). In contrast, there were substantial differences (i.e. 1507 genes) in the transcriptomes of het vs. null NSCs on cuprizone for 3 weeks (Fig. 3, Table 2), consistent with their very distinct responses to demyelination. Many of these genes were downregulated in the nulls in agreement with the role of Gli1 as a transcriptional activator ^49,50^. In the forebrain, Gli1 is expressed by neural stem cells in the SVZ and by mature astrocytes in the parenchyma outside the SVZ excluding the white matter tracts^26,51^. Consistently, the expression of stem cell marker genes (*Sox2, GFAP, Nestin*) were higher than genes expressed in differentiated cells (*APC, Htra1, S100β, PDGFRα, SYNGAP1*) in both *Gli1^HET^* and *Gli1^NULL^* healthy control NSCs.

**Figure 3:**
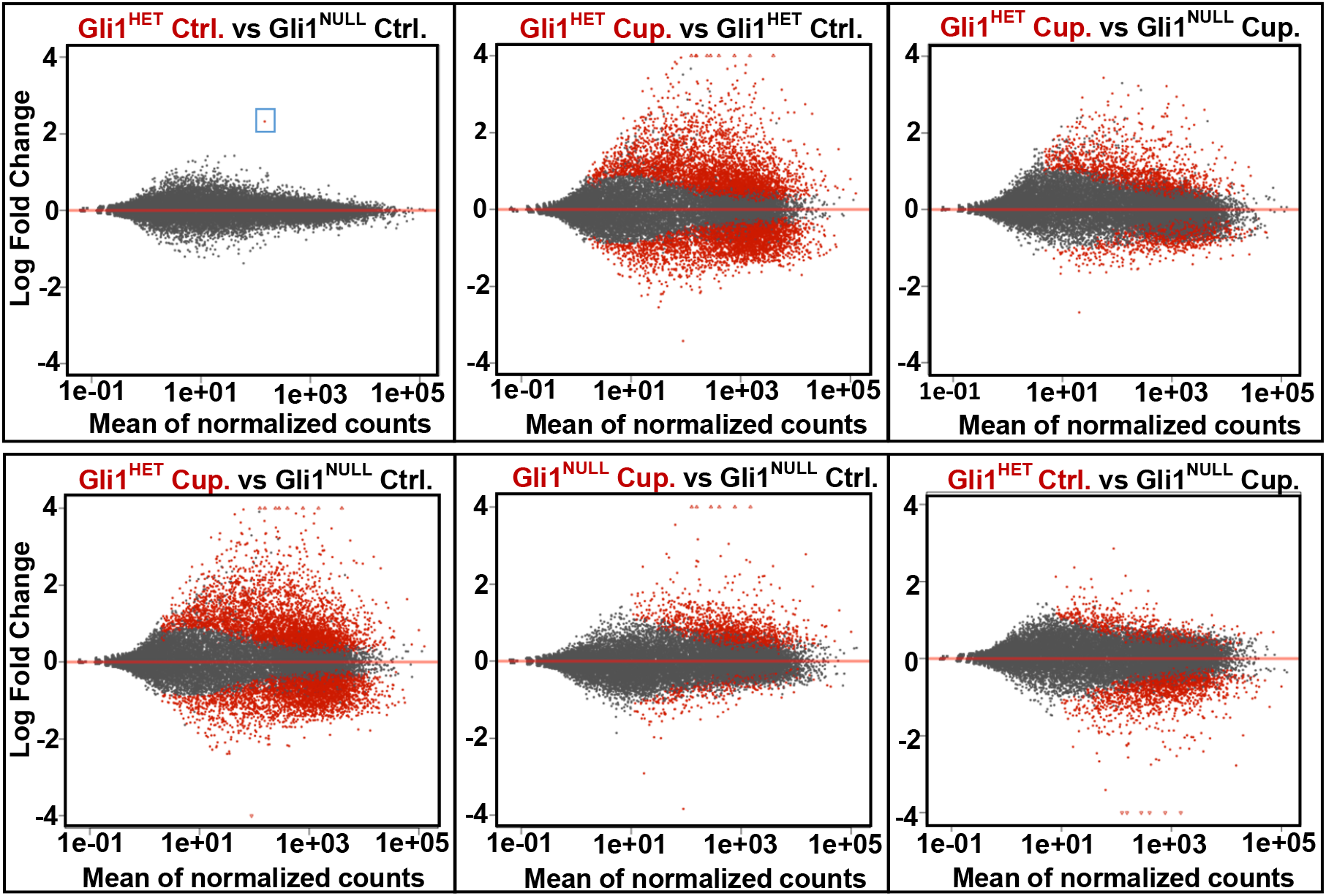
Visualization of differential gene expression in Gli1 het and null NSCs MA plots of the RNAseq data show the differences in expression of genes between the two genotypes on control and cuprizone diets. The blue box highlights the only gene differentially expressed between Gli1^Het^ and Gli1^Null^ NSCs on the control diet groups, corresponding to Gli1 itself. Ctrl.-Control diet group, Cup.-Cuprizone diet group.

**Table 1:** Number of differentially expressed genes. Genes that had an adjusted p-value (False Discovery Rate) of less than 0.05 and fold change of more than 2 or less than 0.5 in the compared groups are quantified.

| Samples | Accession |
| --- | --- |
| Gli1-Het control diet-1 | GSM4956878 |
| Gli1-Het control diet-2 | GSM4956879 |
| Gli1-Het control diet-3 | GSM4956880 |
| Gli1-Het cuprizone diet-1 | GSM4956881 |
| Gli1-Het cuprizone diet-2 | GSM4956882 |
| Gli1-Het cuprizone diet-3 | GSM4956883 |
| Gli1-Null control diet-1 | GSM4956884 |
| Gli1-Null control diet-2 | GSM4956885 |
| Gli1-Null control diet-3 | GSM4956886 |
| Gli1-Null cuprizone diet-1 | GSM4956887 |
| Gli1-Null cuprizone diet-2 | GSM4956888 |
| Gli1-Null cuprizone diet-3 | GSM4956889 |

**Table 2:** NCBI GEO accession numbers of samples Accession number of the samples included in the the NCBI GEO submission GSE162683.

| Comparison | No. of genes:<br>P<0.05 | No. of upregulated<br>genes: P<0.05 | No. of downregulated<br>genes: P<0.05 | No. of genes:<br>P<0.05 & FC>2 | No. of genes:<br>P<0.05 & FC<0.5 |
| --- | --- | --- | --- | --- | --- |
| Gli1<sup>HET</sup> Control vs Gli1<sup>NULL</sup> Control | 1 | 1 | 0 | 1 | 0 |
| Gli1<sup>HET</sup> Cuprizone vs Gli1<sup>HET</sup> Control | 6442 | 3481 | 2961 | 1447 | 525 |
| Gli1<sup>HET</sup> Cuprizone vs Gli1<sup>NULL</sup> Cuprizone | 1507 | 955 | 552 | 500 | 64 |
| Gli1<sup>HET</sup> Cuprizone vs Gli1<sup>NULL</sup> Control | 6447 | 3499 | 2948 | 1482 | 526 |
| Gli1<sup>NULL</sup> Cuprizone vs Gli1<sup>NULL</sup> Control | 1209 | 1015 | 194 | 294 | 34 |
| Gli1<sup>HET</sup> Control vs Gli1<sup>NULL</sup> Cuprizone | 1730 | 436 | 1294 | 86 | 242 |

### Clustering of RNAseq data

A 3D Principle Component Analysis (PCA) plot visualized by Plotly (Fig.4) showed clustering of the samples in the control diet groups from both *Gli1^Het^* and *Gli1^Null^* neural stem cells; similarly the cuprizone diet RNA samples from both genotypes clustered together but away from the control diet groups. We observed an increased variance in the *Gli1^Het^* samples of both control and cuprizone diet groups. However, the heatmaps for the differentially expressed genes (FDR<0.05) showed clustering of *Gli1^Het^* and *Gli1^Null^* samples in the control groups (Fig.5). When we compared the differentially expressed genes (FDR<0.05) between the cuprizone groups, the expression in the *Gli1^Null^* samples appeared similar to the expression in the control groups (Fig.5), suggesting that demyelination results in lesser perturbation of gene expression in *Gli1^Null^* NSCs compared to *Gli1^Het^* NSCs.

**Figure 4:**
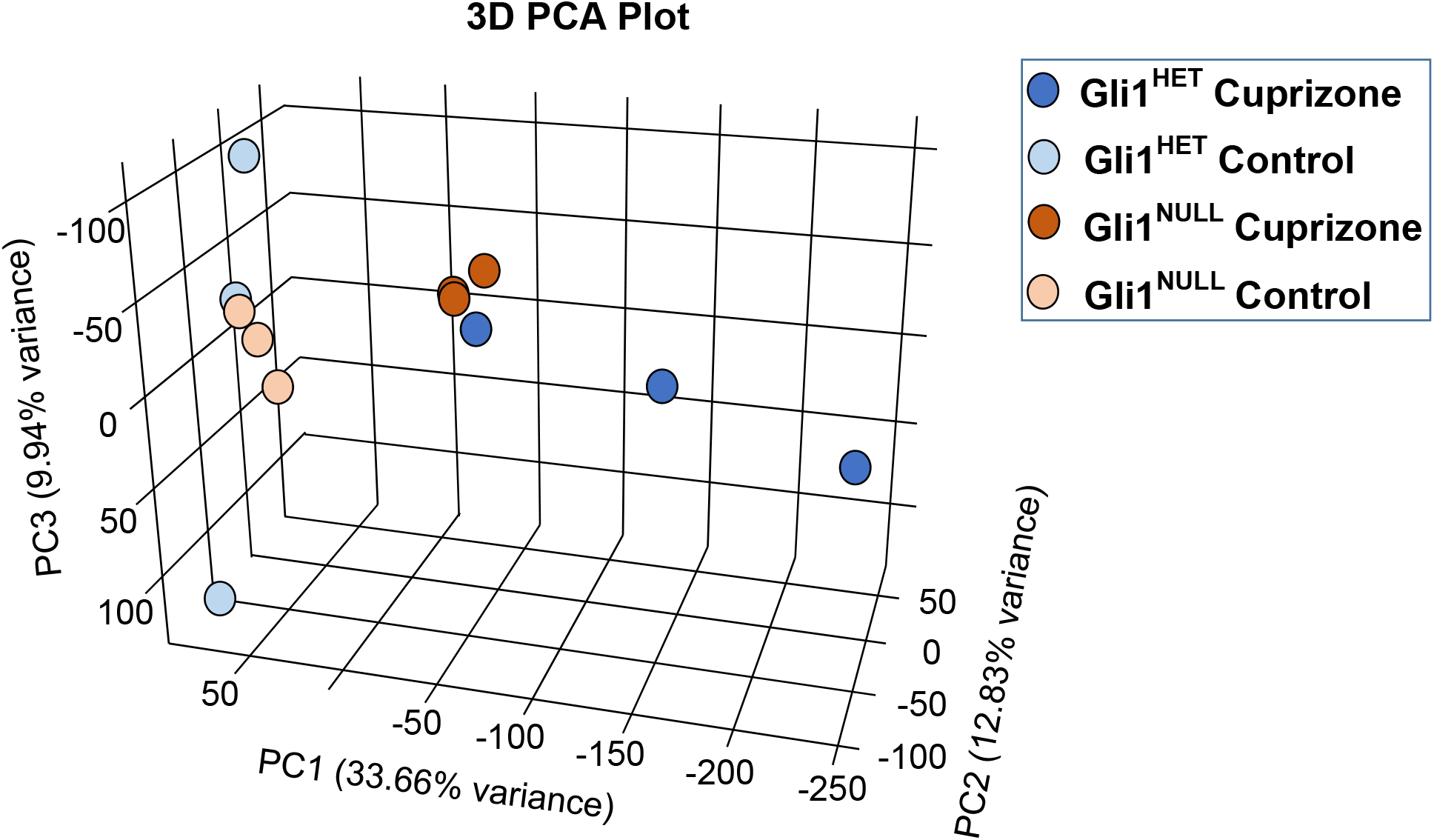
Principle component analysis (PCA) of the sequencing data. A 3D PCA plot shows segregation of the control diet samples from the cuprizone diet samples. The variance is higher in the Gli1^Het^ samples compared to the Gli1^Null^ samples.

**Figure 5:**
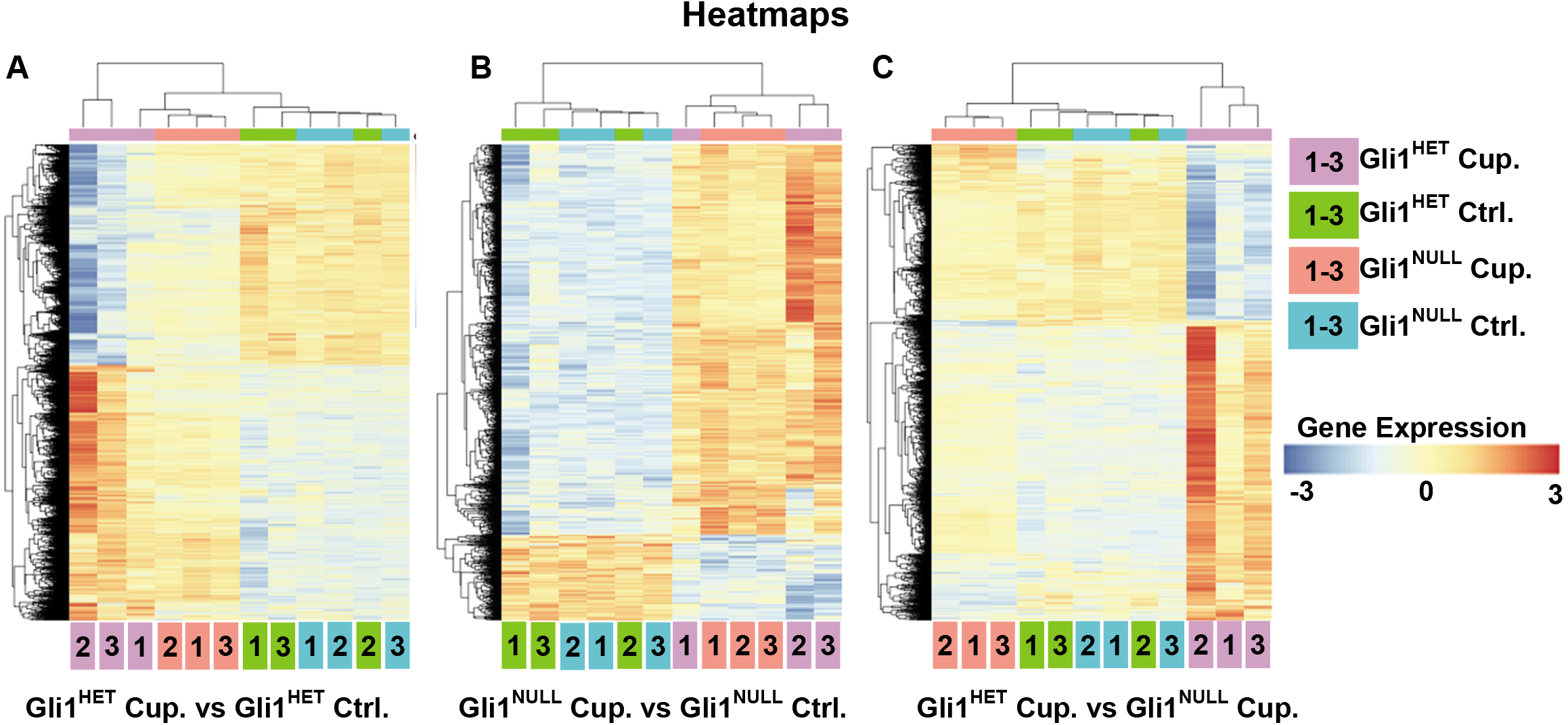
Heatmaps of the differentially expressed genes (FDR<0.05). (A) Comparison of *Gli1^Het^* NSCs following demyelination with cuprizone diet (Cup.) (purple) vs. healthy NSCs on control diet (Ctrl.) (green). (B) Comparison of *Gli1^Null^* NSCs following demyelination with cuprizone diet (Cup.) (red) vs. healthy NSCs on control diet (Ctrl.) (blue). (C) Comparison of the *Gli1^Het^* vs. *Gli1^Null^* NSCs following demyelination with cuprizone diet (Cup.). The sample numbers are mentioned inside the color coded box for each condition, at the bottom of the heatmap. The gene expression levels are color coded with the highest upregulated genes in red and the most down-regulated genes in blue.

### Ingenuity Pathway Analysis

To examine the altered response of *Gli1^Null^* NSCs to demyelination compared to *Gli1^Het^* NSCs, we analyzed 1507 differentially expressed genes (p<0.05), between *Gli1^Het^* and *Gli1^Null^* cuprizone groups with Ingenuity Pathway Analysis (Qiagen, Build version: 448560M) (Table 3). While neurological diseases scored highest amongst the perturbed networks, the cellular functions related to survival and development were amongst the top 5 significant differences between the 2 groups. Consistently, TP53 and TGFβ1 were the most significant upstream regulators and these pathways were predicted to be more active in *Gli1^Het^* NSCs upon demyelination.

**Table 3:** Summary of Ingenuity Pathway Analysis of differentially expressed genes in *Gli1^HET^* cuprizone vs *Gli1^NULL^* cuprizone groups. Top 5 perturbed networks, physiological system functions, molecular and cellular functions and upstream regulators are listed in addition to the top 10 upregulated and downregulated genes in *Gli1^HET^* cuprizone NSCs.

| Networks | Score |
| --- | --- |
| 1. Developmental Disorder, Hereditary Disorder, Neurological Disease | 46 |
| 2. Cellular Assembly and Organization, Cellular Function and Maintenance, Cellular Development | 43 |
| 3. Cell Cycle, Cellular Assembly and Organization, DNA Replication, Recombination, and Repair | 38 |
| 4. Lipid Metabolism, Small Molecule Biochemistry, Auditory Disease | 36 |
| 5. Organ Morphology, Organismal Development, Organismal Injury and Abnormalities | 36 |

| Physiological System Development and Function | p Value | # Genes |
| --- | --- | --- |
| Organismal Survival | 6.94E-07 - 5.57E-25 | 417 |
| Tissue Development | 3.23E-06 - 3.98E-21 | 551 |
| Tissue Morphology | 3.78E-06 - 2.74E-16 | 399 |
| Embryonic Development | 3.23E-06 - 6.79E-15 | 357 |
| Organismal Development | 3.78E-06 - 6.79E-15 | 551 |

| Molecular and Cellular Functions | p Value | # Genes |
| --- | --- | --- |
| Cellular Growth and Proliferation | 3.13E-06 - 8.04E-26 | 645 |
| Cell Morphology | 3.74E-06 - 3.28E-25 | 458 |
| Cell Death and Survival | 3.51E-06 - 1.91E-21 | 515 |
| Cellular Assembly and Organization | 1.41E-06 - 1.04E-17 | 276 |
| Cellular Function and Maintenance | 3.74E-06 - 1.04E-17 | 453 |

| Upstream Regulators | p-value of overlap | Predicted Activation |
| --- | --- | --- |
| TP53 | 9.68E-25 | Activated |
| TGFβ1 | 9.99E-19 | Activated |
| TNF | 1.38E-15 | Activated |
| PDGF BB | 9.79E-15 | Activated |
| KRAS | 1.88E-14 |  |

| Top 10 Upregulated Genes (Log 2-fold expression value) | Top 10 Downregulated Genes (Log 2-fold expression value) |
| --- | --- |
| AHNAK2 (↑3.438) | GJD2 (↓-2.684) |
| CLCF1 (↑3.226) | DENND1C (↓-1.647) |
| ANXA3 (↑3.219) | MAG (↓-1.637) |
| ADM (↑3.027) | ADGRL4 (↓-1.603) |
| GPNMB (↑2.986) | BARD1 (↓-1.558) |
| IGFBP3 (↑2.944) | LRRTM3 (↓-1.553) |
| PLAUR (↑2.923) | GRM1 (↓-1.539) |
| ARHGAP22 (↑2.816) | LOC102634852 (↓-1.501) |
| FLNC (↑2.757) | PGBD5 (↓-1.494) |
| SYTL2 (↑2.722) | Ly6a (↓-1.481) |

### Code Availability

The codes used for aligning the RNAseq reads to the mm10 mouse reference genome with TopHat v2.1.1, quantifying the read counts using HTSeqCount, comparison of the gene expression using DESeq2, and PCA analysis using Plotly 3-D PCA, are available in Figshare^46^.

## ACKNOWLEDGEMENTS

We would like to thank Adriana Hugey and Tenzin Lakham from the NYU Genome Technology Center for performing the RNA sequencing and bioinformatic analysis. This work was supported by grants from NINDS (NS0100867), the New York State DOH (DOH01-STEM5-2016-00305), and the NYU Office of Technology Alliance to JLS.

## AUTHOR CONTRIBUTIONS

JS designed and performed the experiments, and wrote the manuscript. HMS and JJL helped with FACsorting. JLS oversaw the project and edited the manuscript.

## ADDITIONAL INFORMATION

**Correspondence** and request for materials should be addressed to JS and JLS.

## REFERENCES

1 Compston, A. & Coles, A. Multiple sclerosis. Lancet 372, 1502–1517, doi:10.1016/S0140-6736(08)61620-7 (2008).

2 Smith, E. J., Blakemore, W. F. & McDonald, W. I. Central remyelination restores secure conduction. Nature 280, 395–396 (1979).

3 Dubois-Dalcq, M., Ffrench-Constant, C. & Franklin, R. J. Enhancing central nervous system remyelination in multiple sclerosis. Neuron 48, 9–12 (2005).

4 Trapp, B. D. & Nave, K. A. Multiple sclerosis: an immune or neurodegenerative disorder? Annu Rev Neurosci 31, 247–269, doi:10.1146/annurev.neuro.30.051606.094313 (2008).

5 Bunge, M. B., Bunge, R. P. & Ris, H. Ultrastructural study of remyelination in an experimental lesion in adult cat spinal cord. J Biophys Biochem Cytol 10, 67–94 (1961).

6 Lassmann, H., Bruck, W., Lucchinetti, C. & Rodriguez, M. Remyelination in multiple sclerosis. Mult Scler 3, 133–136 (1997).

7 Perier, O. & Gregoire, A. Electron microscopic features of multiple sclerosis lesions. Brain 88, 937–952 (1965).

8 Prineas, J. W., Barnard, R. O., Kwon, E. E., Sharer, L. R. & Cho, E. S. Multiple sclerosis: remyelination of nascent lesions. Annals of neurology 33, 137–151 (1993).

9 Lubetzki, C., Zalc, B., Williams, A., Stadelmann, C. & Stankoff, B. Remyelination in multiple sclerosis: from basic science to clinical translation. Lancet Neurol 19, 678–688, doi:10.1016/S1474-4422(20)30140-X (2020).

10 Franklin, R. J. M. & Ffrench-Constant, C. Regenerating CNS myelin - from mechanisms to experimental medicines. Nat Rev Neurosci 18, 753–769, doi:10.1038/nrn.2017.136 (2017).

11 Menn, B. et al. Origin of oligodendrocytes in the subventricular zone of the adult brain. J Neurosci 26, 7907–7918, doi:10.1523/JNEUROSCI.1299-06.2006 (2006).

12 Nait-Oumesmar, B. et al. Activation of the subventricular zone in multiple sclerosis: evidence for early glial progenitors. Proceedings of the National Academy of Sciences of the United States of America 104, 4694–4699 (2007).

13 Gensert, J. M. & Goldman, J. E. Endogenous progenitors remyelinate demyelinated axons in the adult CNS. Neuron 19, 197–203 (1997).

14 Serwanski, D. R., Rasmussen, A. L., Brunquell, C. B., Perkins, S. S. & Nishiyama, A. Sequential Contribution of Parenchymal and Neural Stem Cell-Derived Oligodendrocyte Precursor Cells toward Remyelination. Neuroglia 1, 91–105, doi:10.3390/neuroglia1010008 (2018).

15 Watanabe, M., Toyama, Y. & Nishiyama, A. Differentiation of proliferated NG2-positive glial progenitor cells in a remyelinating lesion. Journal of Neuroscience Research 69, 826–836, doi:10.1002/jnr.10338 (2002).

16 Zawadzka, M. et al. CNS-resident glial progenitor/stem cells produce Schwann cells as well as oligodendrocytes during repair of CNS demyelination. Cell Stem Cell 6, 578–590, doi:10.1016/j.stem.2010.04.002 (2010).

17 Duncan, I. D. et al. The adult oligodendrocyte can participate in remyelination. Proceedings of the National Academy of Sciences of the United States of America 115, E11807–E11816, doi:10.1073/pnas.1808064115 (2018).

18 Yeung, M. S. Y. et al. Dynamics of oligodendrocyte generation in multiple sclerosis. Nature 566, 538–542, doi:10.1038/s41586-018-0842-3 (2019).

19 Redwine, J. M. & Armstrong, R. C. In vivo proliferation of oligodendrocyte progenitors expressing PDGFalphaR during early remyelination. Journal of neurobiology 37, 413–428 (1998).

20 Sanchez, M. A. & Armstrong, R. C. Postnatal Sonic hedgehog (Shh) responsive cells give rise to oligodendrocyte lineage cells during myelination and in adulthood contribute to remyelination. Exp Neurol 299, 122–136, doi:10.1016/j.expneurol.2017.10.010 (2018).

21 Picard-Riera, N. et al. Experimental autoimmune encephalomyelitis mobilizes neural progenitors from the subventricular zone to undergo oligodendrogenesis in adult mice. Proceedings of the National Academy of Sciences of the United States of America 99, 13211–13216 (2002).

22 Xing, Y. L. et al. Adult neural precursor cells from the subventricular zone contribute significantly to oligodendrocyte regeneration and remyelination. The Journal of neuroscience : the official journal of the Society for Neuroscience 34, 14128–14146, doi:10.1523/JNEUROSCI.3491-13.2014 (2014).

23 Butti, E. et al. Neural stem cells of the subventricular zone contribute to neuroprotection of the corpus callosum after cuprizone-induced demyelination. J Neurosci, doi:10.1523/JNEUROSCI.0227-18.2019 (2019).

24 Doetsch, F., Caille, I., Lim, D. A., Garcia-Verdugo, J. M. & Alvarez-Buylla, A. Subventricular zone astrocytes are neural stem cells in the adult mammalian brain. Cell 97, 703–716 (1999).

25 Chaker, Z., Codega, P. & Doetsch, F. A mosaic world: puzzles revealed by adult neural stem cell heterogeneity. Wiley Interdiscip Rev Dev Biol 5, 640–658, doi:10.1002/wdev.248 (2016).

26 Ahn, S. & Joyner, A. L. In vivo analysis of quiescent adult neural stem cells responding to Sonic hedgehog. Nature 437, 894–897, doi:10.1038/nature03994 (2005).

27 Samanta, J. et al. Inhibition of Gli1 mobilizes endogenous neural stem cells for remyelination. Nature 526, 448–452, doi:10.1038/nature14957 (2015).

28 Namchaiw, P. et al. Temporal and partial inhibition of GLI1 in neural stem cells (NSCs) results in the early maturation of NSC derived oligodendrocytes in vitro. Stem Cell Res Ther 10, 272, doi:10.1186/s13287-019-1374-y (2019).

29 Sanchez, M. A., Sullivan, G. M. & Armstrong, R. C. Genetic detection of Sonic hedgehog (Shh) expression and cellular response in the progression of acute through chronic demyelination and remyelination. Neurobiol Dis 115, 145–156, doi:10.1016/j.nbd.2018.04.003 (2018).

30 Jiayuan, S. et al. Gant61 ameliorates CCl4-induced liver fibrosis by inhibition of Hedgehog signaling activity. Toxicol Appl Pharmacol 387, 114853, doi:10.1016/j.taap.2019.114853 (2020).

31 Gupta, V., Gupta, I., Park, J., Bram, Y. & Schwartz, R. E. Hedgehog Signaling Demarcates a Niche of Fibrogenic Peribiliary Mesenchymal Cells. Gastroenterology 159, 624–638 e629, doi:10.1053/j.gastro.2020.03.075 (2020).

32 Ren, D. et al. Saikosaponin B2 attenuates kidney fibrosis via inhibiting the Hedgehog Pathway. Phytomedicine 67, 153163, doi:10.1016/j.phymed.2019.153163 (2020).

33 Weiskirchen, R., Weiskirchen, S. & Tacke, F. Organ and tissue fibrosis: Molecular signals, cellular mechanisms and translational implications. Mol Aspects Med 65, 2–15, doi:10.1016/j.mam.2018.06.003 (2019).

34 Tsukui, T. et al. Gli signaling pathway modulates fibroblast activation and facilitates scar formation in pulmonary fibrosis. Biochem Biophys Res Commun 514, 684–690, doi:10.1016/j.bbrc.2019.05.011 (2019).

35 Schneider, R. K. et al. Gli1(+) Mesenchymal Stromal Cells Are a Key Driver of Bone Marrow Fibrosis and an Important Cellular Therapeutic Target. Cell Stem Cell 23, 308–309, doi:10.1016/j.stem.2018.07.006 (2018).

36 Kramann, R. & Schneider, R. K. The identification of fibrosis-driving myofibroblast precursors reveals new therapeutic avenues in myelofibrosis. Blood 131, 2111–2119, doi:10.1182/blood-2018-02-834820 (2018).

37 Guerrero-Juarez, C. F. & Plikus, M. V. Gli-fully Halting the Progression of Fibrosis. Cell Stem Cell 20, 735–736, doi:10.1016/j.stem.2017.05.003 (2017).

38 Radecki, D. Z. et al. Relative Levels of Gli1 and Gli2 Determine the Response of Ventral Neural Stem Cells to Demyelination. Stem Cell Reports 15, 1047–1055, doi:10.1016/j.stemcr.2020.10.003 (2020).

39 Matsushima, G. K. & Morell, P. The neurotoxicant, cuprizone, as a model to study demyelination and remyelination in the central nervous system. Brain pathology (Zurich, Switzerland) 11, 107–116 (2001).

40 Caprariello, A. V. et al. Biochemically altered myelin triggers autoimmune demyelination. Proceedings of the National Academy of Sciences of the United States of America 115, 5528–5533, doi:10.1073/pnas.1721115115 (2018).

41 Hanley, O. et al. Parallel Pbx-Dependent Pathways Govern the Coalescence and Fate of Motor Columns. Neuron 91, 1005–1020, doi:https://doi.org/10.1016/j.neuron.2016.07.043 (2016).

42 Strober, W. Trypan Blue Exclusion Test of Cell Viability. Curr Protoc Immunol 111, A3.B.1–A3.B.3, doi:10.1002/0471142735.ima03bs111 (2015).

43 Aken, B. L. et al. The Ensembl gene annotation system. Database (Oxford) 2016, baw093, doi:10.1093/database/baw093 (2016).

44 Love, M. I., Huber, W. & Anders, S. Moderated estimation of fold change and dispersion for RNA-seq data with DESeq2. Genome Biology 15, 550, doi:10.1186/s13059-014-0550-8 (2014).

45 Samanta, J., Salzer, J.L. Transcriptomic analysis of loss of Gli1 in neural stem cells responding to demyelination in the mouse brain. NCBI GEO https://www.ncbi.nlm.nih.gov/geo/query/acc.cgi?acc=GSE162683 (2021).

46 Samanta, J., Salzer, J.L. Transcriptomic analysis of loss of Gli1 in neural stem cells responding to demyelination in the adult mouse brain. figshare https://figshare.com/s/db4525a356708bf43487 (2021).

47 Fleige, S. & Pfaffl, M. W. RNA integrity and the effect on the real-time qRT-PCR performance. Molecular aspects of medicine 27, 126–139, doi:10.1016/j.mam.2005.12.003 (2006).

48 Bai, C. B., Auerbach, W., Lee, J. S., Stephen, D. & Joyner, A. L. Gli2, but not Gli1, is required for initial Shh signaling and ectopic activation of the Shh pathway. Development 129, 4753–4761 (2002).

49 Lipinski, R. J., Gipp, J. J., Zhang, J., Doles, J. D. & Bushman, W. Unique and complimentary activities of the Gli transcription factors in Hedgehog signaling. Exp Cell Res 312, 1925–1938, doi:10.1016/j.yexcr.2006.02.019 (2006).

50 Tolosa, E. J. et al. GLI1/GLI2 functional interplay is required to control Hedgehog/GLI targets gene expression. Biochem J 477, 3131–3145, doi:10.1042/BCJ20200335 (2020).

51 Garcia, A. D., Petrova, R., Eng, L. & Joyner, A. L. Sonic hedgehog regulates discrete populations of astrocytes in the adult mouse forebrain. The Journal of neuroscience : the official journal of the Society for Neuroscience 30, 13597–13608, doi:10.1523/JNEUROSCI.0830-10.2010 (2010).

